# Intermediate Progenitor cells provide a transition between hematopoietic progenitors and their differentiated descendants

**DOI:** 10.1101/2020.10.12.336743

**Authors:** Carrie M. Spratford, Lauren M. Goins, Fangtao Chi, Juliet R. Girard, Savannah N. Macias, Vivien W. Ho, Utpal Banerjee

## Abstract

Genetic and genomic analysis in *Drosophila* suggests that hematopoietic progenitors likely transition into terminal fates via intermediate progenitors (IPs) with some characteristics of either, but perhaps maintaining IP-specific markers. In the past, IPs have not been directly visualized and investigated due to lack of appropriate genetic tools. Here we report a split-*GAL4* construct, *CHIZ-GAL4*, that identifies IPs as cells physically juxtaposed between true progenitors and differentiating hemocytes. IPs comprise a distinct cell type with a unique cell-cycle profile and they remain multipotent for all blood cell fates. Additionally, through their dynamic control of the Notch ligand, Serrate, IPs specify the fate of direct neighbors. The Ras pathway controls the number of IP cells and promotes their transition into differentiating cells. The split-*GAL4* strategy is amenable for adoption in mammalian systems and would be invaluable in assigning trajectories that stem and progenitor populations follow as they develop into mature blood cells.

## Introduction

The transition from a multipotent progenitor into various types of mature, functional cells is a widely studied process in both *Drosophila* and vertebrates. Foundational studies investigating human hematopoiesis revealed the ability of a multipotent hematopoietic stem cell to differentiate into multiple distinct blood cell types (reviewed in: Dzierzak and Speck, 2008; Weissman and Shizuru, 2008). The stem/progenitor and differentiated cell populations are identified and further characterized by expression of unique markers (Coffman and Weissman, 1981; Ikuta and Weissman, 1992; Morrison and Weissman, 1994; Muller-Sieburg et al., 1986; Smith et al., 1991; Uchida and Weissman, 1992). However, the intermediary stage between a stem cell and a differentiated cell is often not well-studied due to a lack of developed tools to target this particular population. *Drosophila* provides an ideal model system with a variety of powerful molecular genetic tools available with which to test and define the function of these intermediate-state cells during the process of hematopoiesis.

Blood cells in *Drosophila* are functionally akin to those derived from mammalian myeloid lineages (reviewed in Evans et al., 2003). As in all invertebrates, *Drosophila* lack lymphoid cells that enable adaptive immunity in vertebrates. The *Drosophila* lymph gland (LG) is the primary site of hematopoiesis during larval development and is made up of multiple paired lobes flanking the dorsal vessel, which functions as the heart (Jung et al., 2005; Mandal et al., 2004). The lymph gland lobes disintegrate during pupariation and the dispersed mature blood cells contribute to the hematopoietic repertoire of the pupa and the adult (Dey et al., 2016; Grigorian et al., 2011).

The anteriorly located lobes are the largest and are referred to as primary lobes that follow a stereotypic pattern of differentiation. Several zones consisting of distinct cell populations have been identified in the primary lobe. The medially located Medullary Zone (MZ) is composed of blood progenitors while the Cortical Zone (CZ) houses three types of mature blood cells (Jung et al., 2005). When present, a cell population termed the Posterior Signaling Center (PSC) functions as a niche and produces a variety of secreted signaling ligands that promote progenitor maintenance (Benmimoun et al., 2015; Lebestky et al., 2003; Mandal et al., 2007; Mondal et al., 2011; Oyallon et al., 2016). The cells of the PSC are defined by their expression of the homeotic gene Antennapedia (Antp) (Mandal et al., 2007).

During first and early second instars, the small primary lobes consist of progenitors that express *domeless* (*dome*) (Jung et al., 2005; Krzemien et al., 2007; Mondal et al., 2011). Hemocyte differentiation initiates at mid-second instar and is marked by *Hemolectin* (*Hml*) and Peroxidasin (Pxn) expression in the developing blood cells (Irving et al., 2005; Jung et al., 2005; Sinenko et al., 2009; Stofanko et al., 2008). Later in the second and third instar larvae, the number of differentiated cells expands forming a distinct CZ. The progenitors populate the MZ and continue to express *dome*. The three mature blood cell types: plasmatocytes, crystal cells, and lamellocytes occupy the CZ (Evans et al., 2009; Jung et al., 2005; Krzemien et al., 2010; Minakhina and Steward, 2010). Mature plasmatocytes are positively identified by the presence of the P1 antigen encoded by the *Nimrod C1 (NimC1)* gene (Kurucz et al., 2007). Crystal cells express Lozenge (Lz), Hindsight (Hnt), and Pro-phenoloxidase (PPO) proteins (Jung et al., 2005; Lebestky et al., 2000; Lebestky et al., 2003; Neyen et al., 2015; Terriente-Felix et al., 2013). Lamellocytes are rarely observed in the lymph gland, but when present they are marked by the L1 antigen encoded by *Atilla* (Honti et al., 2009; Lanot et al., 2001; Markus et al., 2005; Markus et al., 2009; Sorrentino et al., 2002).

A small number of cells residing at the juxtaposition of the MZ and CZ express both *dome* and Pxn but lack mature hemocyte markers, P1 and Lz (Krzemien et al., 2010; Sinenko et al., 2009). This observation suggests a role for these cells in the process of transition from a progenitor to a differentiated fate. Collectively, these cells are referred to as Intermediate Progenitors (IPs) belonging to an Intermediate Zone (IZ) (Krzemien et al., 2010; Oyallon et al., 2016). However, thus far no reporter, enhancer, antibody, or driver exists to specifically identify or genetically alter the intermediate progenitors. For this reason, molecular pathways that regulate maturation of these transitional cells remain unknown. Here we describe the development of a “split-*GAL4*” driver that targets IPs and allows us to monitor and investigate this unique set of transitioning cells. We demonstrate that the IPs are a distinct population of cells that can be increased or reduced in number through genetic manipulation. These cells are multipotent and contribute to all three differentiated blood cell types. These IZ cells have a distinct mitotic and gene expression profile compared to cells of the MZ and CZ.

## Results

### Characterization of the Intermediate Zone cell population of IPs

Using a combination of direct drivers of *domeless* (*dome*^*MESO*^*-GFP*) and *Hemolectin* (*Hml*^*Δ*^*-DsRed*), the IZ cells are seen as an overlapping population at the site of juxtaposition between the MZ and CZ (Figure 1A). In an effort to positively label and manipulate genetic pathways within these intermediate progenitors, we designed a “split-*GAL4*” driver to target the cells with overlapping expression of *dome*^*MESO*^*-GFP* and *Hml*^*Δ*^*-DsRed* (Figure 1B). In these constructs, the *dome*^*MESO*^ enhancer is fused to the p65 activation domain and the *Hml*^*Δ*^ enhancer is used to drive the GAL4 DNA binding domain such that only cells that simultaneously express *dome* and *Hml* drive transgene expression downstream of *UAS* binding sites. For reasons of brevity, we refer to this driver as *CHIZ-GAL4 (<u>C</u>ombined <u>H</u>ematopoietic Intermediate <u>Z</u>one-GAL4). CHIZ-GAL4* is the first identified positive marker for the intermediate zone that reliably labels cells in transition from a progenitor to a mature hemocyte. In this paper we use the terms “Intermediate Progenitors (IPs)”, “intermediate zone (IZ) cells”, and “*CHIZ* cells” interchangeably.

**Figure 1:**
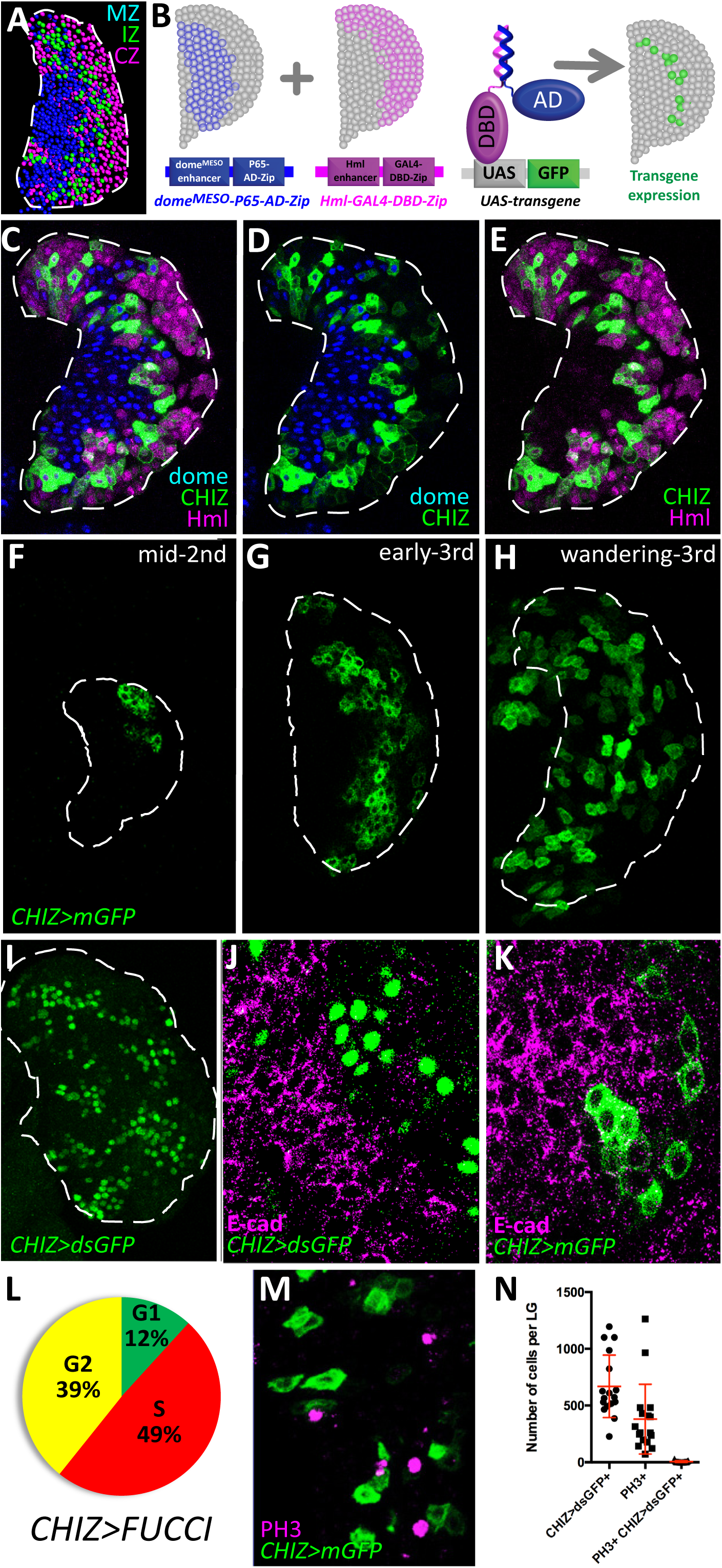
Characterization of intermediate zone cell population. **(A)** Computer rendering of a confocal image of a lymph gland (*dome*^*MESO*^*-GFPnls, Hml*^*Δ*^*-DsRednls*). Nuclei have been pseudo-colored based on endogenous fluorescence. Progenitors in the MZ are labeled by *dome*^*MESO*^*-GFP*, and pseudo-colored blue. Differentiated cells in the CZ are labeled by *Hml*^*Δ*^*-DsRed* and are pseudo-colored magenta. IZ cells identified by an overlap in expression of both *dome*^*MESO*^*-GFP* and *Hml*^*Δ*^*-DsRed* are pseudo-colored green. **(B)** Model depicting the split-*GAL4* components used to create *CHIZ-GAL4*. Shown in blue is the expression of a P65 activation domain (AD) in *dome*^*MESO*^+ cells. Shown in magenta is the DNA binding domain (DBD) of GAL4 which is expressed in *Hml*+ cells. Only the intermediate zone cells with overlapping expression of the AD and DBD express GFP shown in green. **(C-E)** A third instar lymph gland with fluorescently labeled zones (*dome*^*MESO*^*-BFP, Hml*^*Δ*^*-DsRed; CHIZ-GAL4, UAS-mGFP*) shows *CHIZ-GAL4* expression (green) juxtaposed between the MZ (*dome*+, blue) and CZ (*Hml*+, magenta). For clarify, for the same lymph gland shown in **C**, the magenta *Hml* channel is omitted in **D**, and the blue *dome* channel is omitted in **E. (F-H)** Developmental progression of *CHIZ-GAL4* expression (*CHIZ-GAL4, UAS-mGFP*). **(F)** The first appearance of *CHIZ-GAL4* is observed at the distal edge of the mid-second instar lymph gland. **(G)** During early third instar, *CHIZ-GAL4* expression appears in more cells, but with cells at the periphery lacking *CHIZ-GAL4* expression. **(H)** In wandering third instar larvae, *CHIZ-GAL4* expression is dispersed throughout the lymph gland. GFP+ cells seen outside of the dashed line belong to the paired primary lobe. **(I)** Intermediate zone marked with nuclearly localized destabilized GFP. **(J)** E-cadherin protein (magenta) present on the progenitor cell membranes ceases its expression within the IZ cells (green). **(K)** IZ cells (green) directly abut E-cadherin positive cells (magenta). **(L)** Pie chart representing the average percent of *CHIZ-GAL4* expressing cells in primary LG lobes that are in G1 (green), S phase (red), and G2/early M phase (yellow) as assessed by the expression of the *Fly FUCCI* indicator. M phase cannot be separately assessed using *Fly FUCCI*. **(M)** Lack of co-localization of *CHIZ* cells (green) with mitotic marker phospho-histone H3 (magenta). **(N)** Data from *CHIZ>dsGFP* lymph glands stained with PH3 show lack of overlap between IPs and PH3+ cells. Images in **C-E, J, K** and **M** are a single slice of a Z-stack image. Images in **F-I** are a maximum intensity projection of the middle third of the lymph gland of a Z-stack. White dashed lines indicate the edges of lymph gland primary lobe in **A, C-I** as discerned from nuclear staining (not shown).

*CHIZ-GAL4* efficiently marks the IPs when used in conjunction with a short-lived fluorophore such as membrane-GFP (mGFP; Figure 1C), *Fly-FUCCI* (Figure 1--figure supplement 1A) or a rapidly degrading form of GFP (*dsGFP*; Li et al., 1998; Wang et al., 2012) (Figure 1I). In a lymph gland fluorescently marked for MZ and CZ cells, *CHIZ>mGFP* (*CHIZ-GAL4; UAS-mGFP*) faithfully labels cells that express both *dome* and *Hml* and lie at the juxtaposition of the MZ and CZ (Figure 1C-E). Imaging and flow cytometry data show that the IZ comprises 11% (+/- 4.7%, n=115 lymph glands) of total cells in the primary lobes of wandering third instar lymph glands. Long-lived fluorophores such as eGFP are not useful to specifically visualize the transitioning IZ population due to their extended perdurance when driven by *CHIZ-GAL4* (Figure 1—figure supplement 1C).

*CHIZ>mGFP* expression initiates in a small number of cells at the periphery of the lymph gland at mid-second instar (Figure 1F). This timing is also coincident with the onset of differentiation. As larval development progresses into the late second and early third instars, IPs increase in number and intensity to form a band of cells in the middle of the LG (Figure 1G). At the wandering third instar, IPs appear scattered throughout the LG and are notably present in more medial regions compared to earlier stages of development (Figure 1H). E-cadherin (E-cad), which is required for proper progenitor maintenance is prominently expressed in the MZ cells (Gao et al., 2013; Gao et al., 2014; Jung et al., 2005) but its expression ceases immediately prior to the initiation of *CHIZ-GAL4* (Figure 1J, K). These data are consistent with our recent transcriptomic analysis of the lymph gland that suggests lack of E-Cad expression as a characteristic feature of the IZ and that has also identified several gene products that are uniquely representative of the IP population (see accompanying paper; Girard et al., 2020).

We next characterized the cell cycle profile of these transitory cells using the *Fly FUCCI* system (*Fluorescent Ubiquitination Cell Cycle Indicator*) (Zielke et al., 2011). We find that a small percentage of *CHIZ* cells are in G1, while the vast majority is in S and G2 (Figure 1L), a result that is also confirmed by flow cytometric analysis (Figure 1—figure supplement 1A, B). As the *Fly FUCCI* system is unable to distinguish between G2 and early mitotic phases, we sought to measure the occurrence of *CHIZ* cells undergoing mitosis. Surprisingly, *CHIZ* cells are never found to co-localize with phosphorylated Histone H3 (pH3) (Figure 1M, N). Thus, IZ cells can be in every phase of the cell cycle except mitosis. To confirm this unexpected result, we utilized loss of function genotypes in the mitosis-promoting kinase Aurora B (AurB). Loss of this protein is expected to prevent condensation and coupling of chromosomes during mitosis leading to large nuclei with replicated chromosomes (Adams et al., 2001; Giet and Glover, 2001). As expected, expression of *AuroraB-RNAi* in the MZ results in small lymph glands with large nuclei (Figure 1—figure supplement 1D, E). In contrast, *AuroraB-RNAi* expressed in the IPs does not give rise to an observable phenotype (Figure 1—figure supplement 1F, G). We conclude that a mitotic population enters the IZ following cell division but is maintained in a pre-mitotic state until it exits the IZ.

### IPs contribute to all mature hemocyte populations

Under normal conditions, *CHIZ* cells do not express P1 or Hnt (Figure 2A, B), which are markers for mature plasmatocytes and crystal cells, respectively (Kurucz et al., 2003; Terriente-Felix et al., 2013). The IZ cells can be largely eliminated by expression of the pro-apoptotic genes *hid* (*head involution defective)* and *rpr* (*reaper*) (Grether et al., 1995; White et al., 1994) driven by *CHIZ-GAL4*. In this genetic background, the CZ population is greatly reduced (Figure 2C-E) as are the individual numbers of P1+ plasmatocytes and Hnt+ crystal cells (Figure 2F-K). This provides an early indication that the IPs lead to the formation of plasmatocytes and crystal cells. We confirmed this suggestion using iTRACE and G-TRACE lineage tracking constructs (Bosch et al., 2016; Evans et al., 2009) to determine the possible developmental fates of *CHIZ* cells. We find that descendants of *CHIZ* cells are capable of committing to either plasmatocyte or to crystal cell fates (Figure 2L, M). Lamellocytes are not observed under normal conditions, but are induced upon larval injury (Crozatier et al., 2004; Markus et al., 2005; Rizki and Rizki, 1991; Rizki and Rizki, 1992). Post-injury lineage tracing experiments show that IPs can also be fated to become lamellocytes (Figure 2N). Taken together, the antibody staining, lineage tracing, and ablation data show that the IZ cells constitute a transitional population of multipotent progenitors that are capable of contributing to the CZ populations of plasmatocytes, crystal cells, and lamellocytes.

**Figure 2:**
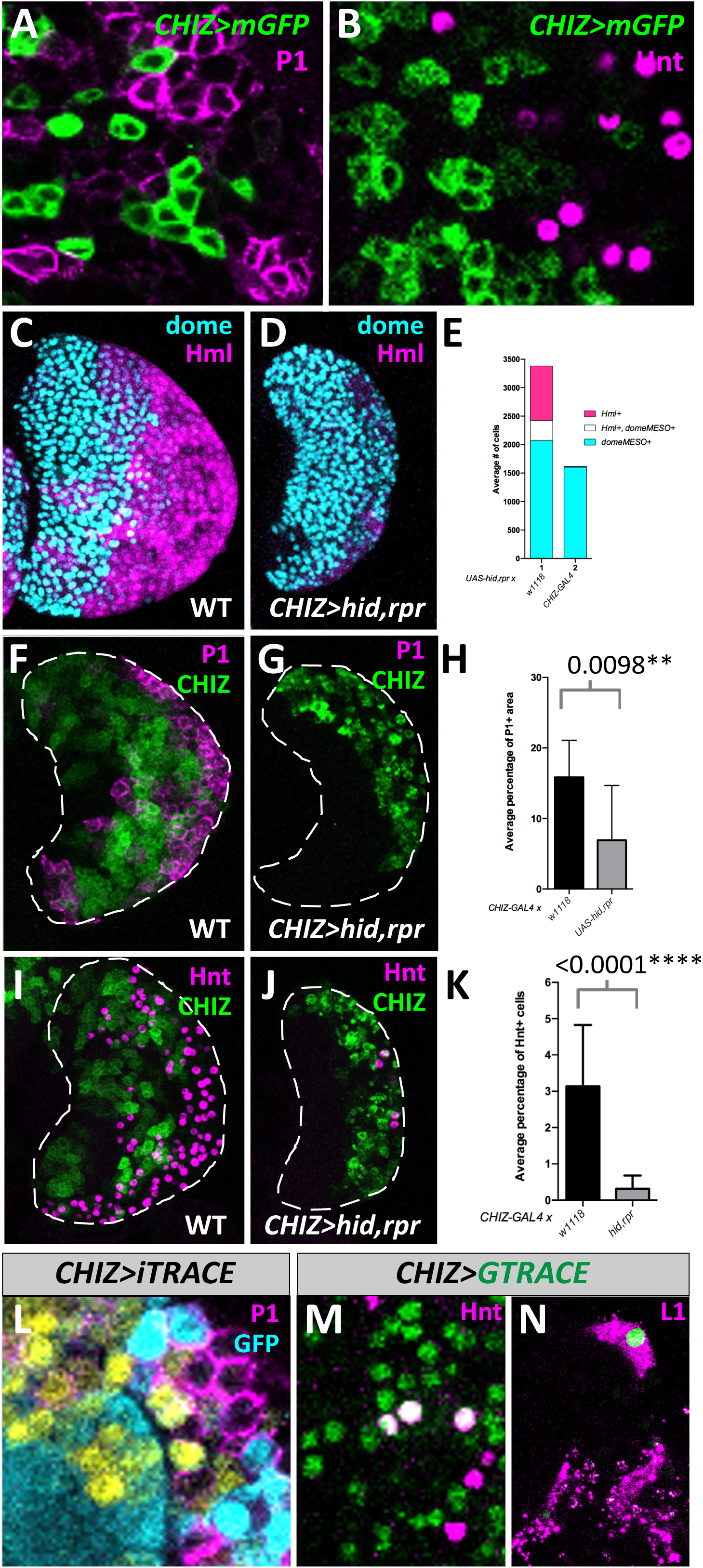
IP cells contribute to all mature hemocyte populations. **(A)** *CHIZ* cells (green) do not co-localize with mature plasmatocytes which stain for P1 (magenta). Instead, *CHIZ* cells are often seen neighboring P1-expressing cells. **(B)** *CHIZ* cells (green) do not stain for Hnt (magenta), a marker for crystal cells. **(C)** A control primary lobe without any GAL4 driver. *dome*+ MZ cells (cyan) and *Hml*+ CZ cells (magenta) (*dome*^*MESO*^*-BFP, Hml*^*Δ*^*-DsRed, UAS-hid,rpr*). **(D)** Apoptosis induced in the IP population leads to a severe decrease in the *Hml*+ (magenta) population compared with *dome*+ (cyan) (*dome*^*MESO*^*-BFP, Hml*^*Δ*^*-DsRed; CHIZ-GAL4, UAS-hid,rpr*). **(E)** Quantitation of data shown in **C, D. (F)** Control showing non-overlap of *CHIZ* cells (green) and P1-expressing cells (magenta) (*CHIZ>mGFP*). **(G)** Genetic ablation of IP cells (green) leads to a reduction in P1-expressing cells (magenta). Also, dying *CHIZ* cells are evident as GFP puncta (green, also seen in **J**) (*CHIZ>mGFP, UAS-hid, rpr*). **(H)** Quantitation of data shown in **F, G. (I)** Control number of Hnt-expressing crystal cells (*CHIZ>mGFP*). **(J)** IP ablation leads to a reduction in crystal cell number (Hnt+, magenta) (*CHIZ>mGFP, UAS-hid, rpr*). **(K)** Quantitation of data shown in **I, J. (L)** *CHIZ* cell descendants (identified by the lack of GFP expression (cyan)) are observed to have P1 antibody staining (magenta). Live expression of *CHIZ-GAL4* is visualized in yellow (*CHIZ-GAL4; UAS-iTRACE*). **(M)** Crystal cells marked by Hnt antibody staining (magenta) can co-localize (white, due to overlap of green and magenta) with cells lineage traced from the *CHIZ* population (green) (*CHIZ-GAL4, UAS-GTRACE*^*LTO*^). **(N)** 24 hours post-injury, cells lineage traced from the *CHIZ* population (green) can be seen expressing L1 (magenta) present in mature lamellocytes (*CHIZ-GAL4; UAS-GTRACE*^*LTO*^). **A** and **B** are single slices from a Z-stack, **L** is a maximum projection stack of 10 slices, **M** and **N** are maximum projection of 3 slices, and **C, D, F, G, I**, and **J** are stacks of the middle third of confocal data. White dashed lines indicate the edges of lymph gland primary lobe in **F, G, I**, and **J**.

### Ras/Raf activity facilitates the IP to hemocyte transition

We next investigated the function of known molecular pathways in the transition between IP and maturing hemocytes. Two major signaling pathways, Ras/Raf and Notch operate during lymph gland development (Crozatier et al., 2004; Dragojlovic-Munther and Martinez-Agosto, 2013; Krzemien et al., 2007; Mondal et al., 2011; Mondal et al., 2014; Mukherjee et al., 2011) but any specific role they might play in the IZ population has not been explored. Later we discuss the role of the Notch pathway in the IPs. Activation of the Ras/Raf/MAPK pathway in *Drosophila* leads to the phosphorylation of Pointed (Pnt) and Yan (Aop), both ETS family proteins that function as downstream transcriptional activator and repressor, respectively (Brunner et al., 1994; Lai and Rubin, 1992; Nusslein-Volhard et al., 1984; O’Neill et al., 1994). Activated forms of Ras or Raf expressed specifically in the IPs causes a reduction in the number of the IPs (Figure 3A-C, F) and reciprocally, inhibition of the Ras pathway increases the IZ population (Figure 3D-F). Concomitantly, we observe an increase in the number of *Hml*+ cells upon activation of the Ras/Raf pathway and a decrease in this population upon loss of function of this pathway (Figure 3G-L). The loss of function phenotype is strikingly apparent when *pnt* expression is knocked down in the *CHIZ* cells (Figure 3J) or when a constitutively active version of *yan* (*Yan*^*ACT*^) (Rebay and Rubin, 1995), expected to block Ras pathway signals, is expressed in the IPs (Figure 3K). This latter result is not phenocopied if the over-expressed version of Yan is wild type (*Yan*^*WT*^, not constitutively activated) (Figure 3L). The phenotypic distinction between *Yan*^*ACT*^ and *Yan*^*WT*^ overexpression further supports the presence of an activated Ras pathway that will cause degradation of the wild-type but not the activated version of Yan. Immunohistochemical localization shows no detectable Yan protein in the IPs (Figure 3M-N), contrary to a previous report, likely due to lack of direct IP markers (Tokusumi et al., 2011). Instead, the Yan protein is detected in crystal cells and *yan-RNAi* expressed in crystal cells eliminates all Yan expression in the lymph gland (Figure 3O and Figure 3--figure supplement 1A-C’). These data characterizing Yan expression are supported by RNAseq results showing high Yan transcript in the crystal cells (Girard et al., 2020). Finally, a subset of *CHIZ* cells express the nuclear form of dp-ERK (active MAPK), but we note that dp-ERK is additionally observed sporadically throughout the lymph gland (Figure 3P-Q). Altogether, these results suggest that cells of the IZ require Ras/Raf signaling to exit the *CHIZ* state and that in the absence of such a signal, the IP cells are held back in their transitional *CHIZ* state.

**Figure 3:**
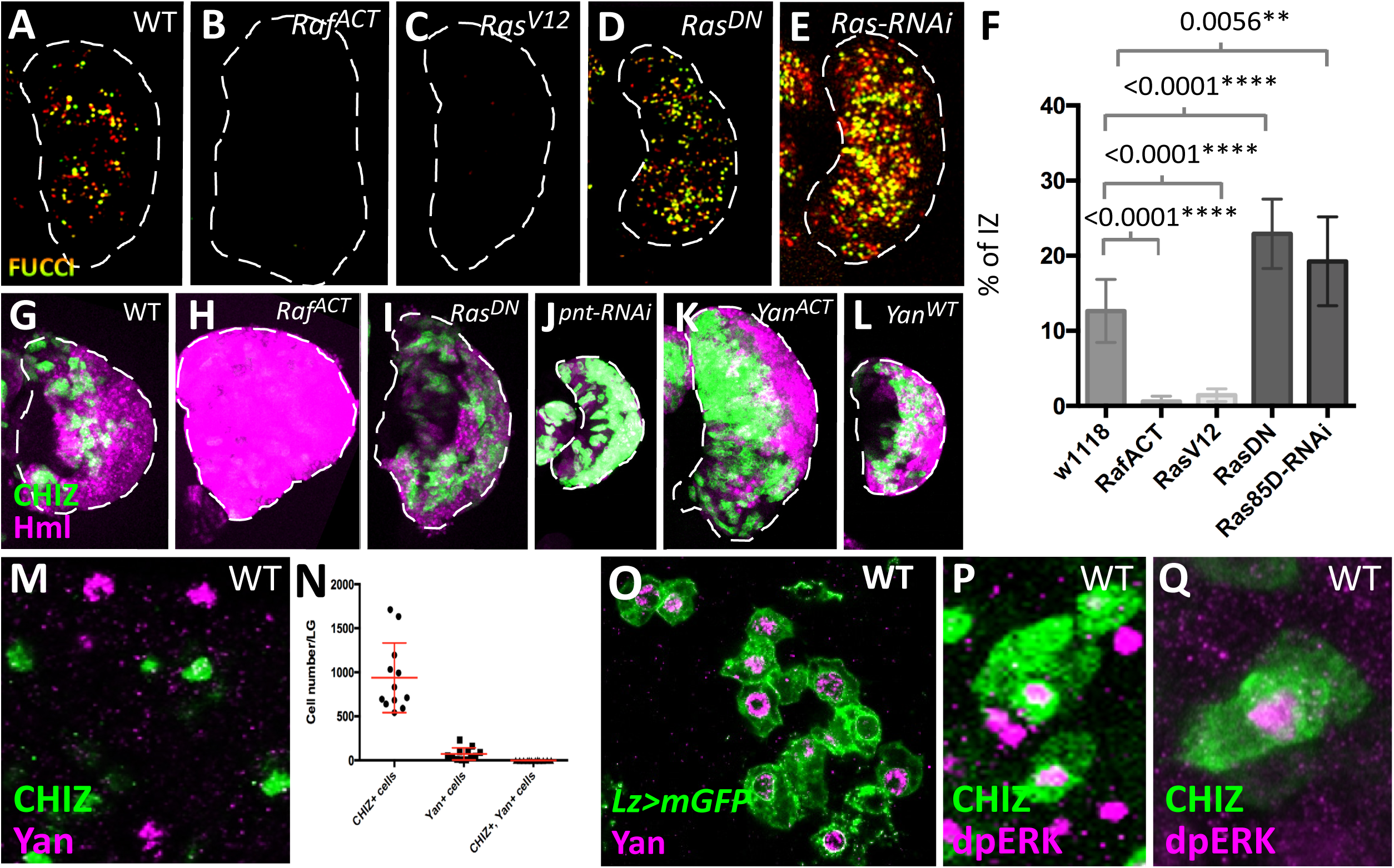
Ras/Raf activity facilitates the IP to hemocyte transition. Images of lymph glands **A-E** and **G-L** are maximum projections of the middle third of a Z-stack. **(A-E)** *CHIZ*+ IZ cells are marked with a nuclear fluorescent marker for quantification purposes (*CHIZ-GAL4, UAS-FUCCI, UAS-X* where X is defined for each panel). **(A)** Control number of IZ cells. **(B)** *UAS-Raf*^*ACT*^ leads to a loss of IZ cells. **(C)** *UAS-Ras*^*V12*^ causes a similar decrease in IZ cells as **B. (D)** An increase in IZ cells is apparent when *CHIZ-GAL4* drives *UAS-Ras*^*DN*^. **(E)** Increased IZ cells are present when expressing *UAS-Ras85D-RNAi* in IZ cells. **(F)** Fraction of *CHIZ*+ cells in lymph glands represented in **A-E. (G-L)** *CHIZ*+ IZ cells are marked in green and *Hml*+ CZ cells are labeled in magenta (*CHIZ-GAL4, UAS-mGFP; Hml*^*Δ*^*-DsRed; UAS-X* where X is defined for each panel). **(G)** Wild type. **(H)** *UAS-Raf*^*ACT*^ causes an extreme expansion of the CZ and loss of IZ. **(I)** *UAS-Ras*^*DN*^ increases the proportion of IZ cells and leads to a decrease in CZ cells. **(J)** *UAS-pnt-RNAi* causes a large increase in proportion of IZ cells and very few CZ cells. **(K)** *UAS-Yan*^*ACT*^ causes an increased IZ and reduced CZ. **(L)** *UAS-Yan*^*WT*^ does not result in a shift in the general proportion of IZ to CZ as seen in **K. (M)** IZ cells (green) do not directly co-localize with nuclear Yan protein (magenta) (*CHIZ>dsGFP*). **(N)** Data from *CHIZ>dsGFP* lymph glands showing lack of any significant overlap between *CHIZ*+ and Yan+ cells. **(O)** Crystal cells (green) express nuclear Yan protein (magenta) (*Lz-GAL4, UAS-mGFP*). **(P, Q)** A subset of IZ cells (green) show nuclear dpERK staining (magenta) (*CHIZ>mGFP*). Images **M, O, P** and **Q** are maximum projection stacks of three slices of a Z-stack.

### IP cells induce crystal cell formation mediated by the Notch pathway

*CHIZ* cells appear at 72 hours after egg lay (hAEL) before the appearance of the first crystal cells (Figure 4A-B). At 96hAEL, a significant number of crystal cells are detected in the immediate vicinity of the *CHIZ* cells with the two cell types separating from each other by 108hAEL (Figure 4A, C-D). This suggests that the close temporal and spatial relationship between these two cell types during early stages of hemocyte differentiation becomes less important as development proceeds to the third instar. Serrate/Notch signaling is important for crystal cell specification in the lymph gland (Mukherjee et al., 2011; Terriente-Felix et al., 2013), therefore we investigated the Notch pathway in the context of IP function. Staining with an antibody raised against the intracellular domain of Notch (Notch^ICD^) shows expression at a low level throughout the lymph gland, with the highest level of staining seen in cells positioned adjacent to *CHIZ* cells (Figure 4E). Interestingly, in the mid-second instar, virtually all of the earliest appearing *CHIZ* cells express high levels of Serrate (Ser). This co-localization of IPs and Ser continues through mid-third instar, although non-overlapping expression is now evident in a fraction of the cells (Figure 4F-G). Importantly, however, knock-down of *Ser* specifically in *CHIZ* cells eliminates Ser protein expression in all cells of the lymph gland (Figure 4H, I). This indicates that all Ser-expressing cells in the lymph gland transition through a *CHIZ*-state at some point in their development and the dynamic pattern of Ser is a reflection of the tight temporal control of its expression. Furthermore, *Ser-RNAi* expressed in *CHIZ* cells causes a significant reduction in the number of crystal cells (Figure 4J). The high Serrate expression is greatly attenuated as the larva matures to the wandering third instar, and at this later stage, there is no correspondence between the residual low Ser-expressing cells and the large number of *CHIZ* cells (Figure 4—figure supplement 1A-A’). RNAseq analysis confirms a high level of Ser expression in IZ cells compared to MZ cells (Girard et al., 2020). We conclude that the IP-specific expression of Ser is dynamic and is responsible for a subset of CZ cells to take on a crystal cell fate.

**Figure 4:**
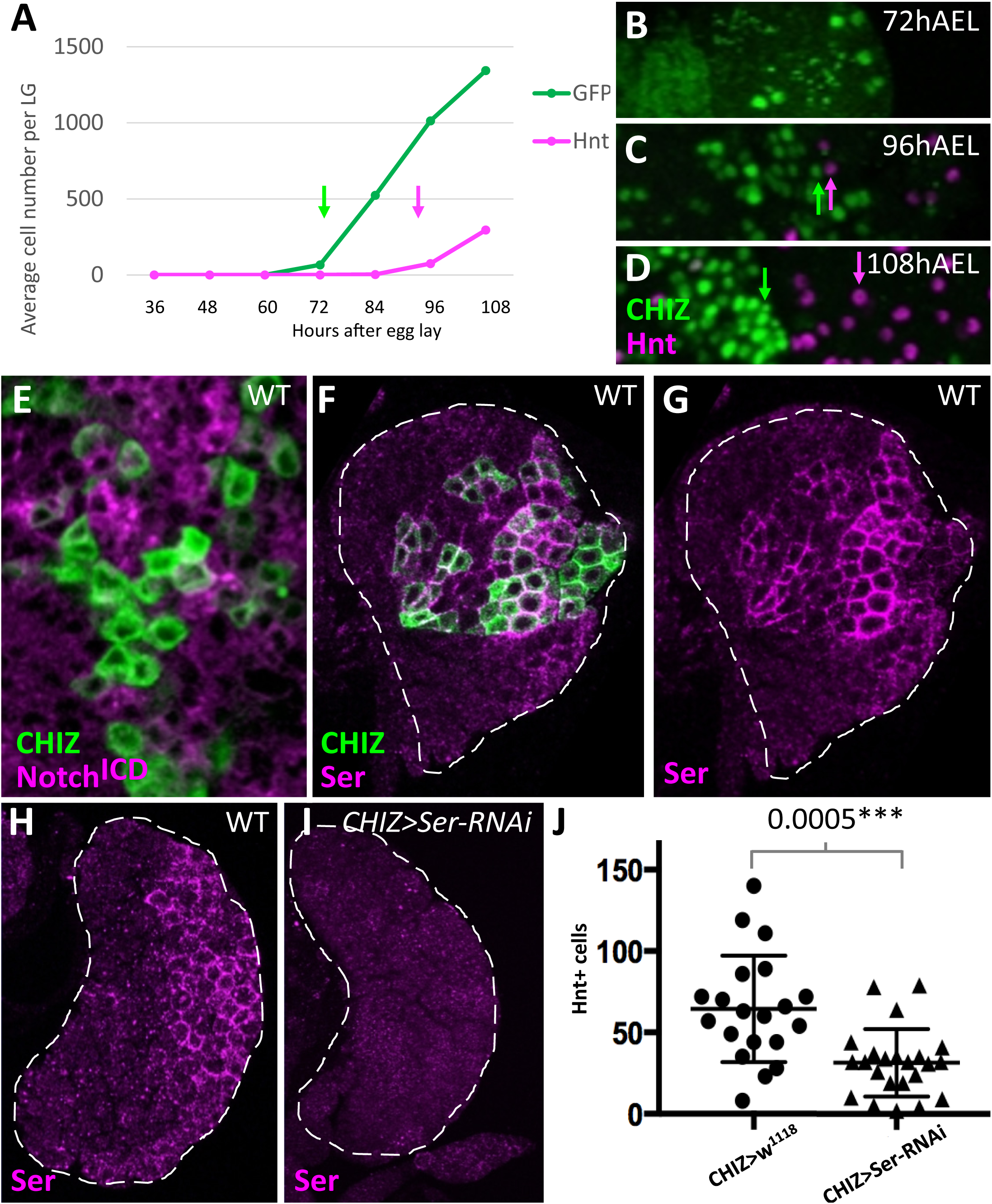
IP cells induce crystal cell formation mediated by the Notch pathway. **(A-D)** Genotype is *CHIZ-GAL4, UAS-dsGFP. CHIZ* cells are green (GFP) and crystal cells are magenta (Hnt). **(A)** Quantification of the raw number of *CHIZ* and crystal cells in lymph glands of developmentally synchronized larvae. The first significant appearance of *CHIZ* cells is at 72 hAEL (green arrow), while the first significant appearance of crystal cells is later, at 96hAEL (magenta arrow). **(B-I)** All images are single slices of confocal data. **(B)** Section from a 72hAEL primary lobe showing the first appearance of *CHIZ* cells. **(C)** At 96hAEL the earliest crystal cells (magenta arrow) are usually seen neighboring *CHIZ* cells (green arrow). **(D)** At 108hAEL wandering third instar, primary lobes have numerous crystal cells (magenta arrow) distant from *CHIZ* cells (green arrow). **(E-G)** Genotype is *CHIZ-GAL4, UAS-mGFP*. **(E)** *CHIZ* cells (green) do not co-localize with high levels of N^ICD^ protein (magenta) observed in neighboring cells. **(F, G)** *CHIZ* cells (green) co-localize with high levels of Serrate protein (magenta) in early third instar. **(H)** Control showing Serrate protein expression (magenta) in early third instar lymph gland. **(I)** Serrate protein (magenta) is absent in early third instar when *Ser-RNAi* is driven by *CHIZ-GAL4* (*CHIZ-GAL4, UAS-Ser-RNAi*). **(J)** The number of crystal cells (Hnt+) per lymph gland decreases when *Ser-RNAi* is expressed in IP cells (*CHIZ-GAL4, UAS-Ser-RNAi*) compared to control (*CHIZ-GAL4*). White dashed lines indicate the edges of lymph gland primary lobe in **F-I**.

## Discussion

Current tools of genome-wide analyses have revealed increased complexities within cell populations that were erstwhile considered homogeneous (Cho et al., 2020; Hernandez et al., 2018; Papalexi and Satija, 2018; Villani et al., 2017; Zeng et al., 2018). Readily available methods for visualization and characterization of cells with such fine distinctions in fate have been more difficult to come by. In this manuscript we used the split-*GAL4* strategy to generate *CHIZ-GAL4* in order to define and describe IPs. Additionally, *CHIZ-GAL4* allows for genetic manipulation of this transitory population that bridges MZ progenitors with the CZ hemocytes. The IPs of the intermediate zone represent a distinct cell type that have some characteristics that are distinct from and others that are similar to the cells of the MZ and CZ. For example, IPs express *dome*, but not E-cadherin, both of which are MZ markers. Similarly, IPs express *Hml*, but not the maturity markers P1 (plasmatocytes) and Hnt (crystal cells). Importantly, we believe IPs to be a unique cell type as their numbers can be expanded or reduced upon genetic manipulation, as shown for example, with modulation of the Ras pathway. Additionally, bulk and single cell RNA-seq data obtained recently in the laboratory identifies several genes that are highly enriched within IPs when compared to their expression in all other cell types in the lymph gland (Girard et al., 2020). In future studies, these will serve well as specific IZ markers and provide further functional relevance for this population.

An important, and perhaps surprising finding of this study is that the IPs are the only cell type in the lymph gland that are found to be in G1, S, and G2 phases of the cell cycle but not M. Several lines of evidence including RNAseq data (Girard et al., 2020) support this conclusion. The MZ cells are also fairly quiescent (Jung et al., 2005; Krzemien et al., 2010; Lebestky et al., 2003), but they are largely held in G2 (Sharma et al., 2019 and L.M.G., J.R.G, and C.M.S. unpublished data), and will undergo mitosis in a limited number of cells. Given that IPs are not restricted to G2, we propose that before entering the IP state, a *dome*+ progenitor is released from G2 and it undergoes mitosis. Subsequently *Hml* is initiated and continues to be expressed as IPs progress through G1, S, and G2. At this point *dome* expression ceases, thus ending the *CHIZ*-state. The *dome*-negative post-*CHIZ* cell likely undergoes a round of mitosis before it progresses to a differentiated state. The IPs are multipotent and contribute to all of the three mature hemocyte populations. Trajectory analysis based on RNA-sequencing is supportive of the above conclusions (Girard et al., 2020). We should note that the data presented in this study do not preclude the possibility that a few of the hemocytes might form by a parallel mechanism that does not involve the IPs.

As in many developmental systems, entry into a proliferative state and fate determination are intimately intertwined and this applies as well to the transition from the IZ to the CZ. We presume that a mitotic event must closely follow exit from the IP state and is linked to differentiation into a hemocyte. We also know that the Ras/Raf pathway is required for exit out of the IP state. In other systems, Ras/Raf activity has largely been associated with proliferation (reviewed in: Bryant et al., 2014; Karnoub and Weinberg, 2008; Lu et al., 2016), but in *Drosophila*, this pathway often governs cell fate determination, as seen for example, during the development of the eye imaginal disc (Flores et al., 2000; Freeman, 1996; Nagaraj and Banerjee, 2007; Simon et al., 1991). Thus, it remains uncertain at the present moment whether Ras/Raf initiates the mitotic process and this allows differentiation signals to be sensed to turn on markers, or if another mechanism controls the entry into mitosis and Ras is responsible for turning off a marker such as *dome*. In a manner similar to that seen in other well-defined developmental situations in *Drosophila*, the Ras/Raf and Notch pathways play dueling roles in the post-*CHIZ* stage of defining cell fate. The IPs express Serrate in a dynamic pattern and induce neighbors to take on a crystal cell fate. The expression of Serrate is downregulated by the mid-third instar, and its restricted spatial and temporal pattern of expression limits crystal cell number. Crystal cells do not have active Ras signaling as established by their expression of the Yan protein. The Ras/Raf signal leads to a plasmatocyte fate, the default pathway seen in the absence of Notch signaling. Upstream events that activate Ras in the IPs will be of great interest for future investigation. It is possible that a canonical ligand-dependent RTK may be involved, however other autonomous molecular inputs could feed into Ras. For example, genes involved in sphingolipid and ceramide signaling are enriched within IPs and are excellent candidates for a Ras-induced metabolic transition of the IPs (Girard et al., 2020).

While this work successfully identifies and manipulates the IPs, the question of why an intermediate state of cells exists still remains. These transitional cells may provide an opportunity for synchronization of cell determination when producing mature hemocytes of different fates such that during normal development, plasmatocytes and crystal cells are created in a stereotypical ratio (Ghosh et al., 2015; Lebestky et al., 2000; Leitao and Sucena, 2015; Tepass et al., 1994). It is also possible that these IPs have unique signaling functions as inferred from their regulation of Serrate expression to induce direct neighbors to take on a crystal cell fate. It is interesting to note that this class of cells can act autonomously as multipotent progenitors while also being non-autonomous inducers of one of the specific blood cell fates. Investigation into the expression of receptors and ligands in IPs will expand our current understanding of the role these cells play in regulating these ratios. We speculate that if proven to function in a similar dual role in other systems, that this presents a mechanism for IPs to maintain a strict balance between the number of true progenitors and each of the determined cell types that they produce.

Additionally, the IP population could exist as a mechanism to regulate the number of cells capable of producing mature hemocytes from the progenitor pool. If all progenitors in the MZ were to directly differentiate into mature hemocytes, a relatively steady pool of progenitors will be difficult to preserve, and the spatio-temporal order of hemocyte specification will not be maintained. The IP population provides a buffer zone that results after a round of mitosis, and a second round of mitosis follows immediately after exit from the *CHIZ* state. An important characteristic of the IP population is its lack of M-phase cells. The coordinated cell cycle within IPs will provide means for rapid immune response. In this context, the IPs could also provide a platform for rebalancing relative numbers of hemocyte types in response to environmental or internally generated stresses such as nutritional, olfactory, redox stress, injury and immune challenge (Cho et al., 2018; Hao and Jin, 2017; Mukherjee et al., 2011; Owusu-Ansah and Banerjee, 2009; Pastor-Pareja et al., 2008; Rizki and Rizki, 1979; Rizki and Rizki, 1992; Shim et al., 2012; Shim et al., 2013; Small et al., 2014).

The experimental strategy used to develop *CHIZ-GAL4* has been successfully adapted for identifying cell types based on the co-expression of other genes in *Drosophila*, particularly in the nervous system (Jenett et al., 2012; Pfeiffer et al., 2008; Pfeiffer et al., 2010). There is nothing about this strategy that is *Drosophila*-specific and one hopes that its most useful application might be to uncover cryptic cell types in the context of the significantly more complex transitions described in mammalian development.

## Author contributions

C.M.S., L.M.G., F.C., J.R.G., S.N.M., and V.W.H. performed experiments. C.M.S., L.M.G., J.R.G., S.N.M., and V.W.H. analyzed data. C.M.S., L.M.G., and U.B. contributed to conceptualization of the project and writing the manuscript. U.B. supervised the project and provided funding.

## Acknowledgements

We would like to thank the members of the Banerjee Lab for their comments and suggestions that improved this project. We thank Gerald Rubin, Nancy Fossett, Istvan Ando, Ken Irvine, Katja Brüeckner, and Brian McCabe for gifts of molecular constructs, antibodies, reagents, protocols, and *Drosophila* stocks that furthered this research. We thank the UCLA Broad Stem Cell Research Center (BSCRC), the BSCRC Flow Cytometry Core, the Department of Molecular, Cell, and Developmental Biology BSCRC Core Facilities in Microscopy, the UCLA *Drosophila* Media Facility, and the Eli and Edythe Broad Center of Regenerative Medicine and Stem Cell Research for providing equipment and services essential to this research. U.B. is supported by National Institutes of Health grants R01 HL-067395 and R01 CA-217608; C.M.S. by Ruth L. Kirschstein National Research Service Award number T32HL863458; L.M.G. by Ruth L. Kirschstein Institutional National Research Service Award number T32CA009056; F.C. by the China Scholarship Council Award and California Institute for Regenerative Medicine Pre-doctoral Fellowship; and J.R.G. by Ruth L. Kirschstein National Research Service Award number T32HL69766 and Institutional Research and Academic Career Development Award number K12GM106996.

## Materials and Methods

### Drosophila stocks and husbandry

The following *Drosophila* stocks were utilized for this study: w^*1118*^ (U. Banerjee), *Hml*^*Δ*^*-DsRed*.*nls* (Katja Brüeckner), *dome*^*MESO*^*-GFP*.*nls, Hml*^*Δ*^*-DsRed*.*nls/CyO* (U. Banerjee), *dome*^*MESO*^*-GAL4-AD, Hml*^*Δ*^*-GAL4-DBD* (a.k.a. *CHIZ-GAL4*, developed in Banerjee Lab for this paper, see below), *UAS-mGFP* (II) (U. Banerjee), *CHIZ-GAL4, UAS-mGFP* (U. Banerjee), *dome*^*MESO*^*-BFP, Hml*^*Δ*^*-DsRed, Hh-GFP/FM7* (U. Banerjee), *UAS-dsGFP* (II) (Brian McCabe), *UAS-FUCCI* (BL55722), *UAS-2xEGFP* (U. Banerjee), *dome*^*MESO*^*-GAL4* (U. Banerjee), *AuroraB-RNAi* (BL28691), *UAS-iTRACE* (BL66387), *UAS-GTRACE*^*LTO*^ (U. Banerjee), *dome*^*MESO*^*-BFP, Hml*^*Δ*^*-DsRed, Hh-GFP, UAS-hid, rpr/FM7* (U. Banerjee), *UAS-hid, rpr* (U. Banerjee), *CHIZ-GAL4, UAS-mGFP; Hml*^*Δ*^*-DsRed* (U. Banerjee), *UAS-Raf*^*ACT*^ (BL2033), *UAS-Ras*^*V12*^ (U. Banerjee), *UAS-Ras*^*DN*^ (U. Banerjee), *UAS-Ras85d-RNAi* (BL29319), *UAS-pnt-RNAi* (BL31936), *UAS-Yan*^*ACT*^ (BL5789), *UAS-Yan*^*WT*^ (BL5790), *Lz-GAL4, UAS-mGFP* (U. Banerjee), *UAS-Yan-RNAi* (BL34909), *UAS-Yan-RNAi* (BL35404), and *Ser-RNAi* (U. Banerjee). All stocks were maintained at room temperature or 18°C. All genetic crosses, with the exclusion of the staged larval time course experiment described below, were raised at 29°C for maximum GAL4-UAS efficiency. All flies were raised on standard *Drosophila* fly food with a recipe containing dextrose, corn meal, and yeast.

### Development of *CHIZ-GAL4* driver

The *dome*^*MESO*^ enhancer (*dM-*Forward primer: 5’-CACCCGTCTACCGCGATTCCAAGCACATCCG-3’; *dM*-Reverse primer: 5’-GGATCCAAAATACCCGATGTAAAATCG-3’), *Hml*^*Δ*^ enhancer (*Hml*^*Δ*^-Forward primer: 5’-CACCGGTACCCAAAAGTTATTTCTG-3’, and *Hml*^*Δ*^-Reverse primer: 5’-GTTTAATTGTATACACAGGAAAATC-3’) were amplified from *Drosophila* genomic DNA and ligated into the pENTR™/D-TOPO™ vector (Invitrogen: Cat#K240020) for Gateway cloning. Each entry vector was ligated into the pBPp65ADZpUw (Addgene 26234) and pBPZpGAL4DBDUw (Addgene 26233) destination vectors using the LR ligase (Invitrogen: Cat# 11791020) to generate the desired vectors (*dome*^*MESO*^*-p65-AD*, and *Hml*^*Δ*^*-pGAL4-DBD*). These vectors were sent to BestGene Inc for microinjection. Transgenic flies were generated by PhiC31 integrase-mediated site-specific transgenesis. The *Hml*^*Δ*^*-pGAL4-DBD* was integrated into the 51C locus of the *Drosophila* genome (Injection Stock: 24482NF), while the *dome*^*MESO*^*-p65-AD* was integrated into the 58A locus (Injection Stock: 24484NF). The transgenic *Drosophila* lines were crossed to generate *dome*^*MESO*^*-p65-AD, Hml*^*Δ*^*-pGAL4-DBD* stable lines through homologous recombination.

### Lymph gland dissection and immunohistochemistry

Larval head complexes were dissected on a silicon dissecting dish in chilled 1xPBS. Head complexes including mouth hooks, eye-antennal discs, brain and ventral nerve cord, and lymph glands were immersed in fixation solution (4% formaldehyde in 1xPBS) for 25 minutes. After fixation, samples were washed three times for 10 minutes in lymph gland wash buffer (0.4% Triton in 1xPBS). Samples were incubated in 10%NGS in 1xPBS blocking solution for 10-30 minutes then incubated in primary antibody overnight at 4°C. Samples were washed in lymph gland wash buffer, then incubated in secondary antibody for 2-4 hours at room temperature. After washing off the secondary antibody in lymph gland wash buffer, ToPro dye (Invitrogen) was incorporated at a 1:1000 concentration for 7-10 minutes to visualize nuclei of tissues. After a final wash step, samples were immersed in Vectashield anti-fade mounting media, placed on a glass slide, and lymph glands were isolated from the head complexes and mounted. Samples were covered with a glass coverslip which was sealed with clear nail polish. Slides were stored at 4°C until imaged.

Primary antibodies used in this study include rabbitαGFP (1:100), ratαEcad (1:20, DSHB #DCAD2), rabbitαPH3 (1:1000, Cell Signaling #97015), mouseαP1 (1:100, Istvan Ando), mouseαHnt (1:200, DSHB #1G9), mouseαNotch^ICD^ (1:100, DSHB #C19.9C6), ratαSerrate (1:1000, Ken Irvine), mouseαYan (1:100, DSHB #8B12H9), rabbitαdpERK (1:100, Cell Signaling #4370), and αL1 (1:10, Istvan Ando). ToPro-3 (1:1000, Invitrogen). Secondary antibodies used in this study were purchased from Invitrogen and include: donkeyαmouse AlexaFluor405, donkeyαmouse AlexaFluor488, donkeyαmouse AlexaFluor555, donkeyαmouse Alexa-Fluor633, donkeyαrabbit AlexaFluor488, donkeyαrabbit AlexaFluor555, donkeyαrat AlexaFluor555, and donkeyαrat Cy3, and donkeyαmouse Cy3 from Jackson Scientific. Secondary antibodies were used at a 1:100-1:2000 dilution dependent on the strength of the primary antibody.

### Staged larval lymph gland dissections

For data collected presented in Figure 4A-D, larvae were synchronized within an hour of each other in 12-hour phases. 100-200 mated flies (*CHIZ-GAL4* x *UAS-dsGFP*) were maintained in collection chambers at 25°C and allowed to lay embryos on plates containing ethyl acetate (EA) media. After a 12-hour collection period, new EA plates were provided to the adults in collection chambers. The embryos on the old EA plates were incubated at 25°C for 24 hours. After this incubation time, hatched larvae were cleared from the plate using a paintbrush and the remaining unhatched embryos were incubated for 1 hour at 25°C. After 1 hour, newly hatched larvae were transferred with a paintbrush to a fresh vial of food. Five larvae were placed in each vial. Vials were incubated at 25°C until samples from all time points were dissected and processed for immunohistochemistry on the same day.

### Microscopy and Image Processing

All samples were imaged using a Zeiss LSM-880 confocal microscope using a Z-stack technique with 1.88uM slice thickness. Images were processed using ImageJ. Unless otherwise noted in the Figure Legends, images of lymph glands are a maximum intensity projection of the stack of the middle third of the samples.

### Data Quantification

All quantifications were performed using Imaris data analysis software by Bitplane to quantify Z-stacks of entire lymph glands. Briefly, lymph glands were contoured and fluorescent channels were masked to restrict quantifications to both primary lobes. To label and count nuclei, a spots filter was applied based on ToPro DNA dye incorporation. The DNA+ spots were then filtered against additional fluorescent channels to quantify specific cell types including *CHIZ>dsGFP*+, PH3+, or Hnt+. *FUCCI*+ cells, *CHIZ*+ PH3+ cells, and *CHIZ*+ Yan+ cells were identified by positively filtering for additional fluorophores. Percent of the lymph gland occupied by these particular cell types was determined by dividing the number of cells of interest by the total number of nuclei per lymph gland, then multiplying by 100. When quantifying the volume of the lymph gland and volume of P1+ fluorescence for data presented in Figure 2H, the surfaces filter was first applied based on ToPro DNA dye incorporation and then extended to fill in the volume of both primary lobes. A second volume measurement was made using the surfaces filter for P1+ fluorescence. The percent of the lymph gland occupied by P1+ fluorescence was calculated by dividing the volume of P1+ fluorescence by the total lymph gland volume, then multiplying by 100. All p-values presented represent unpaired student T-tests to determine statistical significance.

### Flow cytometry

*CHIZ-GAL4, UAS-FUCCI* lymph glands were dissected in 1XMDSS (Modified Dissecting Saline Solution: 9.9 mM HEPES-KOH, 137 mM NaCl, 5.4 mM KCL, 0.17 mM NaH_2_PO_4_, 0.22 mM KH_2_PO_4_, 3.3 mM Glucose, 43.8 mM Sucrose, pH 7.4) and immediately submerged in Schneider’s S2 media in a glass watch glass on ice. Isolated lymph glands were washed once with 1XMDSS. 1xMDSS was removed. 200uL of heat activated Papain solution (100 units/mL) was added to lymph glands which were then moved to an Eppendorf tube. Samples were covered in foil and incubated in Papain solution while shaking at 25° for 15 minutes. During incubation, tubes were removed twice to pipette up and down to break up tissue. Papain solution was inactivated by addition of 500uL cold S2 media. Tissue was centrifuged at 3000rpm for 5 minutes. Supernatant was removed and 1mL of 1% formaldehyde was added to the cell pellet. Cells were shaken in fixative at 4° for 30 minutes. Cells were spun down at 3000rpm for 5 minutes and supernatant was removed. Cell nuclei were labeled by incubating pellet at room temperature for 30 minutes in NucBlue live cell stain Ready Probes Reagent (Invitrogen, Hoechst33342 Special Formulation). Sample was transferred to a round-bottom polystyrene tube and samples were run through a BD LSRII FACS analyzer. Gates for cell fluorescence were standardized using single fluorophore controls. This experiment was replicated five times using 50-85 lymph glands per round.

---

**Figure 1—figure supplement 1:**
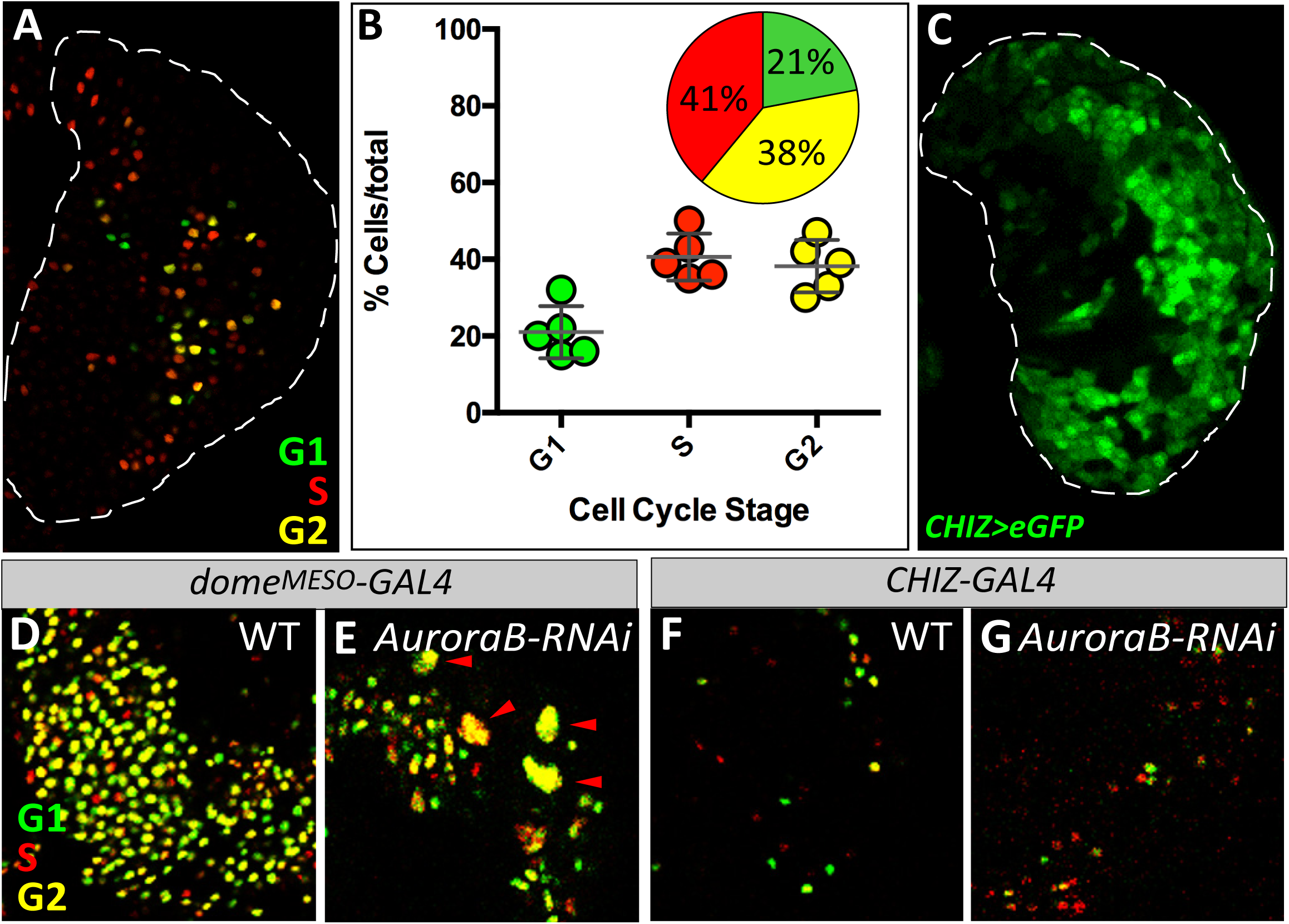
**(A)** IPs can be in G1 (green), S (red), or G2 (yellow) phases of the cell cycle (*CHIZ-GAL4; UAS-FUCCI*). **(B)** Flow cytometric analysis of IPs indicates the majority of IPs are distributed equally between S (red) and G2 (yellow) with a smaller percent of cells in G1 (green) (*CHIZ-GAL4; UAS-FUCCI*). **(C)** Extended perdurance of strong and long-lived fluorophores such as eGFP (green) do not properly represent the specificity of *CHIZ-GAL4* expression. Such fluorophores perdure into the CZ region and fail to follow the transitory nature of the IPs (*CHIZ-GAL4; UAS-2xeGFP*). **(D-G)** Nuclear size-based assay for M-phase cells **(D)** Nuclei of progenitors marked by a cell cycle indicator (*dome*^*MESO*^*-GAL4, UAS-FUCCI*). **(E)** *dome*+ nuclei attempting to enter M-phase at the edge of the MZ become enlarged in size when mitosis is blocked by loss of *AuroraB* (red arrowheads) (*dome*^*MESO*^*-GAL4; UAS-FUCCI, UAS-AuroraB-RNAi*). **(F)** Nuclei of IP cells marked by a cell cycle indicator (*CHIZ-GAL4; UAS-FUCCI*). **(G)** Due to lack of M-phase cells, nuclear size of IPs is not affected upon loss of *AuroraB* (*CHIZ-GAL4; UAS-FUCCI, UAS-AuroraB-RNAi*). White dashed lines indicate the edges of lymph gland primary lobe in **A**, and **C**.

**Figure 3—figure supplement 1:**
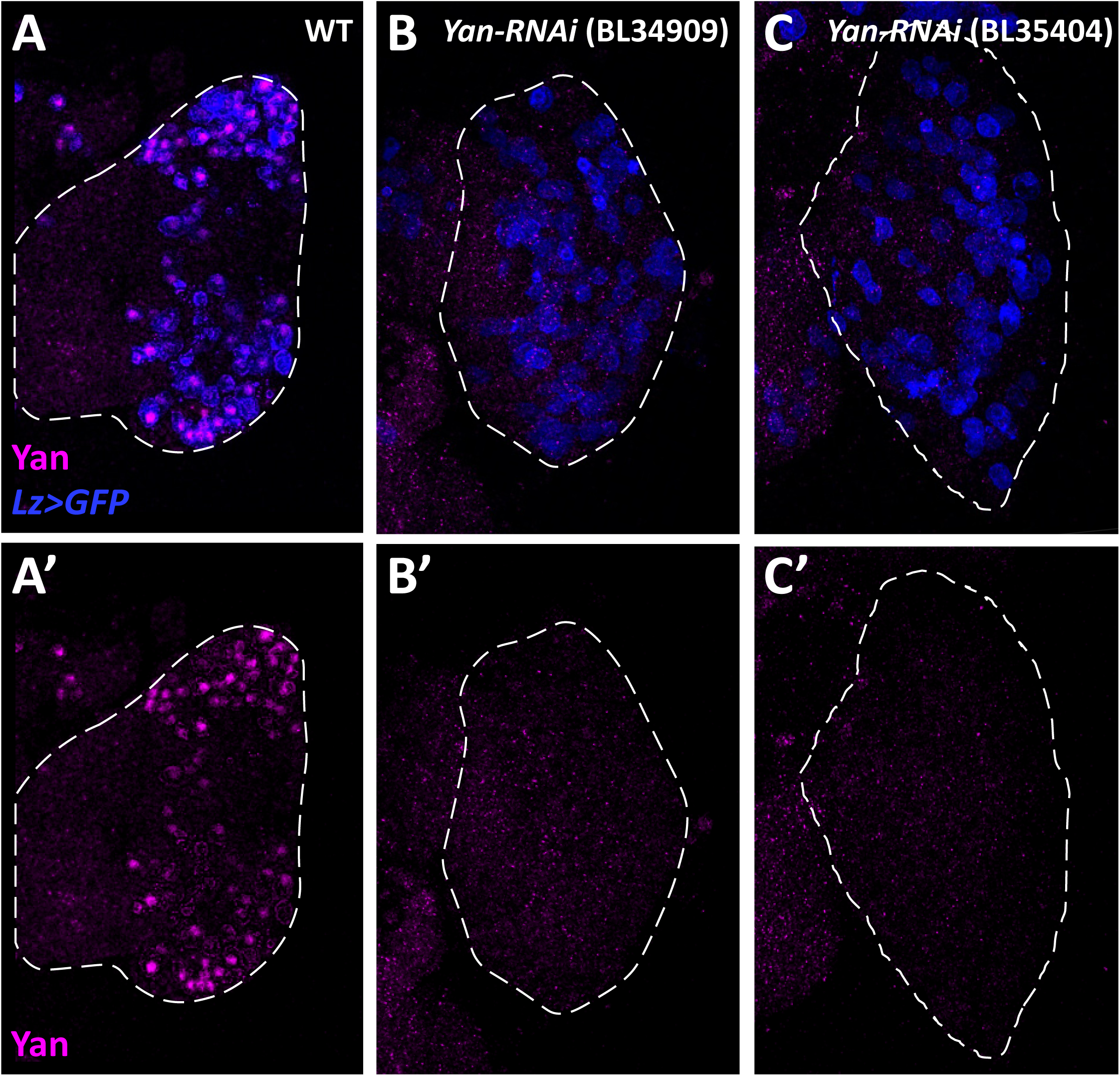
**(A, A’)** Control showing Yan staining (magenta) colocalized with crystal cells (blue) throughout the primary lobe. **(B-C’)** Expression of two separate *Yan-RNAi* constructs, BL34909 **(B**,**B’)** and BL35404 **(C**,**C’)** driven by *Lz-GAL4*. Lz+ crystal cells form **(B, C)**, even though they are devoid of nuclear Yan accumulation **(B’, C’)**.

**Figure 4—figure supplement 1:**
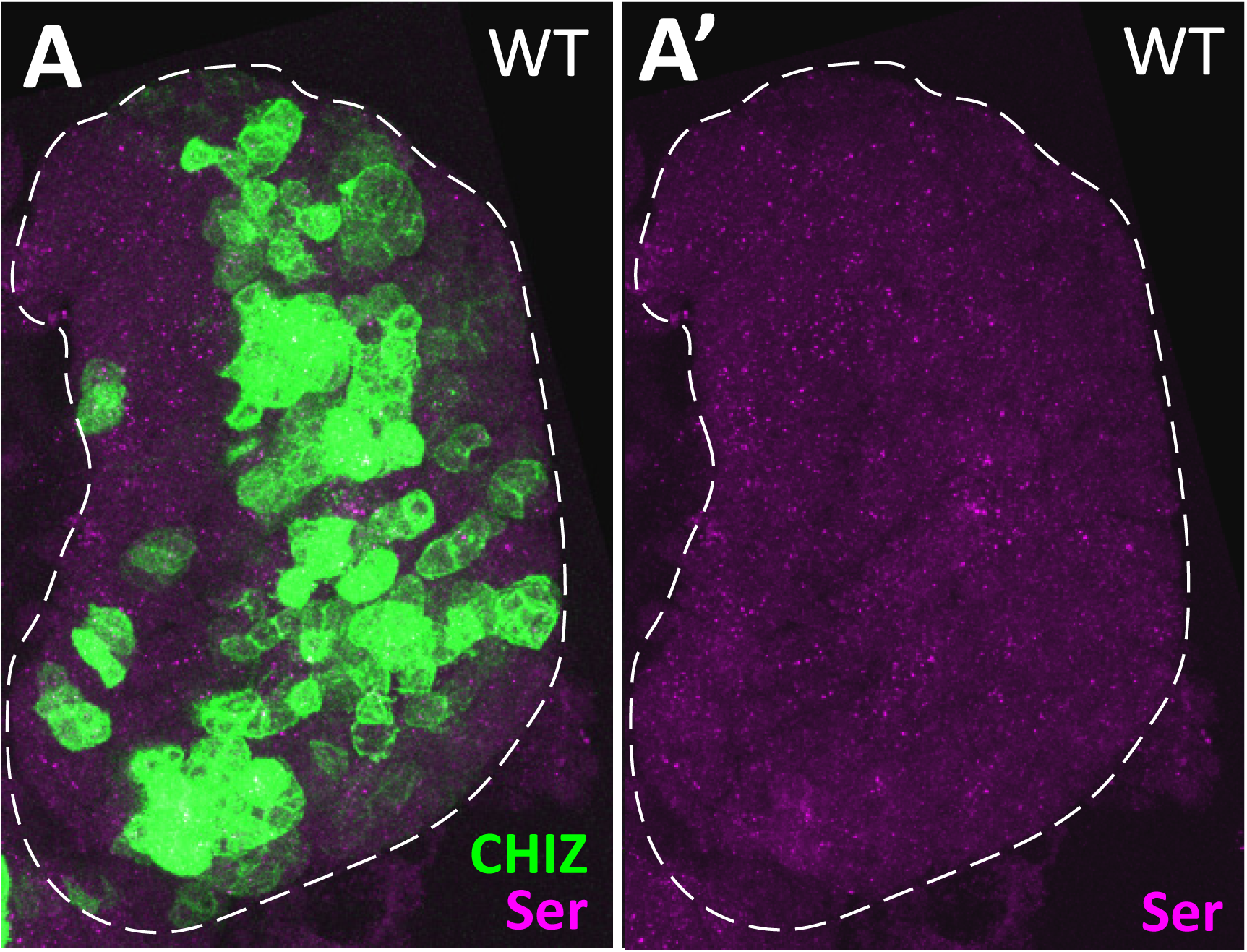
**(A, B)** Control wandering late third instar lymph gland shows Serrate staining is virtually absent (magenta) and this no longer correlates with *CHIZ* cells (green). Images are a maximum projection of the middle third of a confocal Z-stack.

